# *Bacillus* adaptation to *Pseudomonas* secondary metabolites enhances its root competitiveness

**DOI:** 10.64898/2026.07.04.736374

**Authors:** Guillaume Balleux, Marco Zarattini, Adrien Anckaert, Lennard van Buren, Silvia Ribeiro Monteiro, Sébastien Rigali, Marc Ongena

## Abstract

*Bacillus velezensis* is a widely used plant growth-promoting rhizobacterium whose effectiveness under natural conditions is strongly influenced by interactions with surrounding microorganisms. While bacterial secondary metabolites are known to shape these interactions, little is known about their long-term evolutionary consequences. Here, we show that repeated exposure of *B. velezensis* GA1 to secondary metabolites produced by the competing rhizobacterium *Pseudomonas sessilinigenes* CMR12a drives the emergence of an adapted subpopulation with enhanced ecological fitness. Multi-omics analyses revealed extensive metabolomic and transcriptional changes associated with altered growth dynamics, sporulation, motility, and biofilm formation. Importantly, the evolved variant exhibited improved tomato root colonization and reduced the abundance of the competing *Pseudomonas* strain *in planta*. Together, our results demonstrate that prolonged exposure to diffusible bacterial metabolites can drive rapid adaptive diversification in rhizosphere-associated bacteria and highlight the importance of long-term interbacterial interactions in shaping the outcome of plant microbiome assembly and biocontrol performance.

## Introduction

The rhizosphere is the narrow region of soil influenced by plant roots [1]. This soil area represents a dynamic environment densely inhabited by microorganisms, among which bacteria, and to a lesser extent fungi, are the most abundant species [2]. The rhizosphere microbiome considerably influences plant development, health, and resilience, through host-microbe interactions that can be beneficial, commensal or pathogenic [3,4]. Microbe-microbe interspecies interactions ranging from cooperative mutualism to competitive antagonism also reshape the functionalities and composition of plant-associated microbial communities and therefore also influence plant fitness and physiology.

Recent advances in “omics” and analytical technologies have greatly improved our ability to characterize these interactions at multiple levels of resolution. Accumulating evidence shows that bacteria exhibit pronounced physiological plasticity, by modulating the production of secondary/specialized metabolites (SM), transcriptional responses, and phenotypic traits in response to neighbouring microorganisms [5–13]. Such responses can profoundly alter bacterial behaviour, affecting traits involved in resource acquisition, motility, biofilm formation, and root colonisation. Consequently, microbial interactions not only shape community composition but also influence the ability of individual species to establish and persist in the rhizosphere. Understanding these processes is particularly important in the context of microbial inoculants, whose efficacy in promoting plant health often remains inconsistent under field conditions.

The rhizosphere thus represents a highly competitive niche, where keystone bacterial taxa commonly associated with plant roots act as strong competitors for nutrients, colonization sites, and plant-derived resources [14,15]. As a result, interactions between these species frequently lead to antagonistic outcomes, although their metabolite exchanges and adaptive responses may promote coexistence [16]. Such interspecies interactions have been extensively studied in model systems between keystone genera such as *Bacillus* and *Pseudomonas* [16]. They can be neutral [6,17,18] or even beneficial [13], but are also predominantly antagonistic, ranging from amensalism [19] to direct competition [20]. In many cases, chemical exchanges involving different types of SM, such as cyclic lipopeptides, play a central role in shaping the dynamics of these interspecies interactions investigated over short timescales [6,21–23].

In addition to short timescale interactions, recent works deploying the so-called experimental evolution (EE) have revealed that sustained interspecies interactions can impose selective pressure driving adaptive changes in key developmental traits [24,25]. To date, prolonged microbial interactions have largely been studied in co-culture systems involving close physical association, such as adjacent microcolonies and mixed biofilms [26–28]. However, the outcomes of long-term evolutionary adaptation of a rhizobacterium being repeatedly exposed to diffusible SM from neighbouring competitors remains largely understudied.

In this work, we explored the adaptive responses of *Bacillus velezensis* GA1 upon repeated exposure to *Pseudomonas*-derived SM in a contact-less experimental setting. Our data show that chemical communication drives adaptive diversification in *B. velezensis*, affecting both developmental and competitive traits.

## Material and methods

### Strains and media used

All the strains and the species used in this study are listed in Table S1. All bacteria were cultured overnight in shacking Lysogeny Broth (LB) medium (10 g L^−1^ NaCl, 10 g L^−1^ tryptone and 5 g L^−1^ yeast extract at pH = 7). Bacteria were washed three times with physiological water (9 g L^-1^ NaCl) before experimental manipulation. For the evolutional experiments, *B. velezensis* GA1 was grown at 30 °C in half diluted root exudate-mimicking (Re ½) medium (for composition, see Nihorimbere et al., 2012) [29]. Supernatants from *Pseudomonas* strains were collected 48 h cultivation in Re ½ medium, under continuous agitation. Bacterial cultures were filtered (0.22 µm) to obtain the respective cell-free supernatants (CFS).

### GA1.Evo GFP-tagged construction

*B. velezensis* GA1 GFP-tagged was constructed following the method described by Hoff et al., 2021 [30]. Briefly, the GA1 *amyE* fragment was cloned into the pGEM-T Easy vector whereas a *cat*-gfp cassette, bearing a chloramphenicol resistance gene (*cat*) and a promoter lacking *gfpmut3.1* gene, was amplified with KasI restriction sites. The pGEM-T *amyE* plasmid and the *cat*-gfp cassette were digested with KasI (New England BioLabs) and ligated to obtain the pGEM-T *amyE*up-cat-gfp-*amyE*dw construct. Two fragments were first amplified from the pGEM-T *amyE*up-*cat*-gfp-*amyE*dw plasmid: one containing the upstream *amyE* homologous region and the *cat* gene, and the second containing the *gfpmut3.1* gene and the downstream *amyE* homologous region. These two fragments were fused by PCR to generate the complete *cat-*gfp*-amyE* cassette. *B*. *velezensis* GA1 transformation was performed following the protocol developed by Hoff et al., 2021 [30]

### Whole genome sequencing and RNA sequencing

For the genome sequencing, genomic DNA was extracted and purified from *B*. *velezensis* GA1 using the Wizard Genomic DNA purification DNA (Promega) and whole genome sequencing was performed at the Novogene platform (Munich, Germany) by Illumina NovaSeq with paired-end 150 bp reads, following the provider’s standard workflow. DNA quality and quantity were assessed prior to library preparation. Variant calling was performed using BCFtools v.1.19 [31] to identify single nucleotide polymorphisms (SNPs) and insertions/deletions (indels), applying a minimum sequencing depth threshold of 10 reads.

For RNA sequencing (RNA-seq), total RNA was extracted from *B. velezensis* GA1 samples using the NucleoSpin RNA kit (Macherey-Nagel). Total RNAs were quantified using a Nanodrop instrument (ThermoFisher). RNA-seq libraries were prepared and sequenced by Novogene genomics platform (Munich, Germany) using an Illumina NovaSeq platform with a paired-end 150 bp read configuration according to the provider’s standard workflow. Transcriptome data was analyzed using the Seq2science RNA-seq workflow [32]. In short, paired-end reads were trimmed with fastp [33], aligned with STAR [34] to the *B*. *velezensis* GA1 reference genome (CP046386.1), and quantified with Salmon [35]. Differential expression analysis was done with PyDESeq2 [36].

### Secondary metabolites analyses

SM quantification was performed by ultra-high-performance liquid chromatography-mass spectrometry (UPLC-MS) using an Agilent 1290 Infinity II coupled with a mass detector (Jet Stream ESI-Q-TOF 6530) in positive mode. The parameters used were settled up as follows: capillary voltage of 3.5 kV, nebulizer pressure of 35 lb/in2, drying gas of 8 L/min, drying gas temperature of 300 °C, flow rate of sheath gas of 11 L/min, sheath gas temperature of 350°C, fragmentor voltage of 175 V, skimmer voltage of 65 V, and octapole radiofrequency of 750 V. Accurate mass spectra were recorded in the m/z range of 100–1,700. For each analysis, a C18 Acquity UPLC ethylene bridged hybrid column (2.1 mm × 50 mm × 1.7 µm; Waters Corporation, Milford, MA, USA) was used and heated at 40 °C. The injection volume for each sample was 5 µL. Concerning the mobile phase, milliQ water (solvent A) and acetonitrile (solvent B), both acidified with 0.1%v/v of formic acid, were used with a constant flow rate of 0.6 mL/min. The mobile phase gradient used started at 10 % B, kept at this ratio for one minute before rising to 100 % B in 10 min. Solvent B was then kept at 100 % for 3,5 min before coming back to the starting conditions. Data collection and analysis were performed with MassHunter Workstation (version 10.0) and processed with MZmine 3 [37], following parameters described in Table S2.

### Mobility and biofilm formation assays

To assess the swarming and the swimming activities, 5 µL of bacterial culture (OD_600nm_ = 0.1) was inoculated at the centre of Re½ soft agar (7 g L^-1^ of agar) medium and swim agar (3 g L^-1^ agar). Bacterial spreading was imaged through a CoolPix camera (Nikon) and colony areas were quantified using ImageJ software.

Total biofilm quantification was evaluated by crystal violet (CV) staining. Two hundreds µL of bacterial strains (OD_600nm_ = 0.1) were inoculated in a 96-well microplate. Plates were incubated at 30 °C for 24 h without shaking before washing steps with physiological water to remove culture medium and planktonic cells. Biofilms were washed with physiological water to remove culture medium and planktonic cells. Biofilm pellicles were stained with 0.1% (v/v) CV for 15 minutes. The remaining dye was removed by three consecutive washes with physiological water. Violet biofilms were then dissolved with 30% (v/v) acetic acid before measuring the absorbance at 595 nm. Finally, pictures of the biofilm formed were taker with a stereomicroscope (Nikon SMZ1270) equipped with a Nikon DS-Qi2 monochrome microscope camera.

### Total cells and spores counting

Cell-population and spore productions from *Bacillus* strains were quantified by counting. Briefly, bacterial strains (OD_600nm_ = 0.1) were inoculated in 12-wells microplates containing 2 mL of Re ½ medium. Samples were collected after 4, 8, 12, 16 and 20 h of cultivation. At each time point, the entire culture was collected and sonicated for 1 minute to disrupt the biofilm and avoid bias during dilution steps. Following sonication, 500 µL of the culture was incubated at 80 °C for 20 minutes to kill vegetative cells. For both total cell counts and spore counts, serial dilutions were performed. Diluted samples were plated on LB agar plates (14 g L⁻¹ agar), which were then incubated at 30 °C for 16 h. Quantification was performed in three biological replicates, each with four technical replicates.

### In vitro confrontation with Pseudomonas

Confrontation between GA1 strains and *P. sessilinigenes* CMR12a have been performed on Re ½ solid medium (addition of 14 g L^-1^ of agar). Bacteria were grown overnight in liquid Re ½ medium before being washed three times with physiological water (9 g L^-1^ NaCl). Two µL of bacterial suspension (OD_600nm_ = 0.1) was applied at a 5 mm distance from each other and plates incubated at 30 °C for 24 h. Pictures were taken using a CoolPix camera (NIKKOR ×60, as described before).

### *In planta* colonization and confrontation

For *in planta* studies, tomato seeds (*Solanum lycopersicum* var. Moneymaker) were sterilized with ethanol 80 % (v/v), bleach 50 % (v/v) and rinsed at least three time with sterile water before plating in Petri dishes (1 seed per plate) containing Murashige and Skoog (MS) salt and 1 % agar. Seeds were stratified for three days at 4 °C and incubated at 22 °C under a 16/8 h day/night cycle with constant light during photoperiod [30]. Bacteria were inoculated at the root apex of tomato plantlets and roots colonisation comparative studies performed with different GA1 strains (3 µL, OD_600nm_ = 0.1), *P. sessilinigenes* CMR12a (3 µL, OD_600nm_ = 0.1) (mono-inoculation) or a mix of GA1 and CMR12a cells (95:5 ration) (co-inoculation). Following bacterial inoculation, tomato plantlets were growth at 22 °C under the same night/day cycle for three days. Microscopy imaging was performed three days of inoculation with a Nikon Ti2-E inverted microscope (Japan) equipped with a Nikon DS-Qi2 monochrome microscope camera and a ×20/0.45 NA S Plan Fluor objective lens (Nikon, Switzerland). GFP-tagged *Bacillus* and *P*. *sessilinigenes* carrying a mCherry construct were visualized by epifluorescence microscopy. Colonization rates were determined by collecting and weighing plant material and detaching bacteria by vortexing for 5 min in physiological water (9 g L^-1^ NaCl). Cells counting was performed by serial dilution on solid LB (14 g L^-1^ agar) 24 h after incubation at 30 °C for CMR12a and at 37 °C for the GA1 cells.

Regarding the SM production, both for GA1 and CMR12a *in planta* conditions, once the roots used for the colonisation analysis, MS solid area around the plantlets were collected and placed at -20 °C for 24 h. After that, the agar was thawed at room temperature and centrifuged. The supernatants were then collected and filtered (0.22 µm pore size filters) before UPLC-MS analysis.

### Statistical analysis

GraphPad PRISM 8 software was used for statistical analyses. ANOVA or Student paired T-test analysis were performed. The groups that differed significantly from each other, at α = 0.05, were labelled with stars (*).

## Results

### *Long-term exposure to Pseudomonas* metabolites leads to adaptive response in *B. velezensis* evolved sub-population

Previously, it was reported that short-term exposure of *B*. *velezensis* to SM produced by some *Pseudomonas* species triggered marked phenotypic and metabolic changes [6,11,38], with the strongest responses observed upon exposure to compounds secreted by *P. sessilinigenes* or the closely related *Pseudomonas protegens* [6,39–41]. In this work, we first evaluated the potential of cell-free supernatant (CFS) extracts obtained from two additional *Pseudomonas* species (*Pseudomonas simiae* WCS417, *Pseudomonas defensor* WCS374) in addition to *P. sessilinigenes* CMR12a to trigger SM production in *B. velezensis* GA1 in a single batch liquid culture, which we refer as short-term interaction experiments. Upon addition of CFS at subinhibitory dose (4 % v/v), we observed that they were all inducing a *B*. *velezensis* metabolite response after 8h of cultivation (Figure S1). Indeed, upon *Pseudomonas* CFS sensing, *B. velezensis* selectively boosted the production of the cyclic lipopeptide (CLP) surfactin (only with CMR12a and WCS374 CFS), the polyketides bacillaene and dihydrobacillaene and the siderophore bacillibactin, indicating a broad-spectrum response to various *Pseudomonas* competitors. However, among the three CFS tested, the most consistent boost in surfactin, bacillaene and dihydrobacillaene synthesis was measured in response to the extract prepared from *P*. *sessilinigenes* CMR12a. Therefore, we selected this strain for further EE experiments designed to test the effect of prolonged or long-term interaction.

In this set-up, *B. velezensi*s GA1 was grown in a root exudate mimicking medium [29] under static conditions allowing both SM analysis and additional phenotyping. GA1 wild type (hereafter defined as GA1.WT) was cultivated for 13 consecutive serial passages in the presence of CMR12a CFS (hereafter called as adapted lineages, GA1.Ad) or in the absence of it (GA1.Ctrl lineages) (Figure 1a). After each passage, cells from both lineages were washed and part of the cell’s population was either conserved for further characterization or directly used to inoculate culture for the next passage.

**Figure 1:**
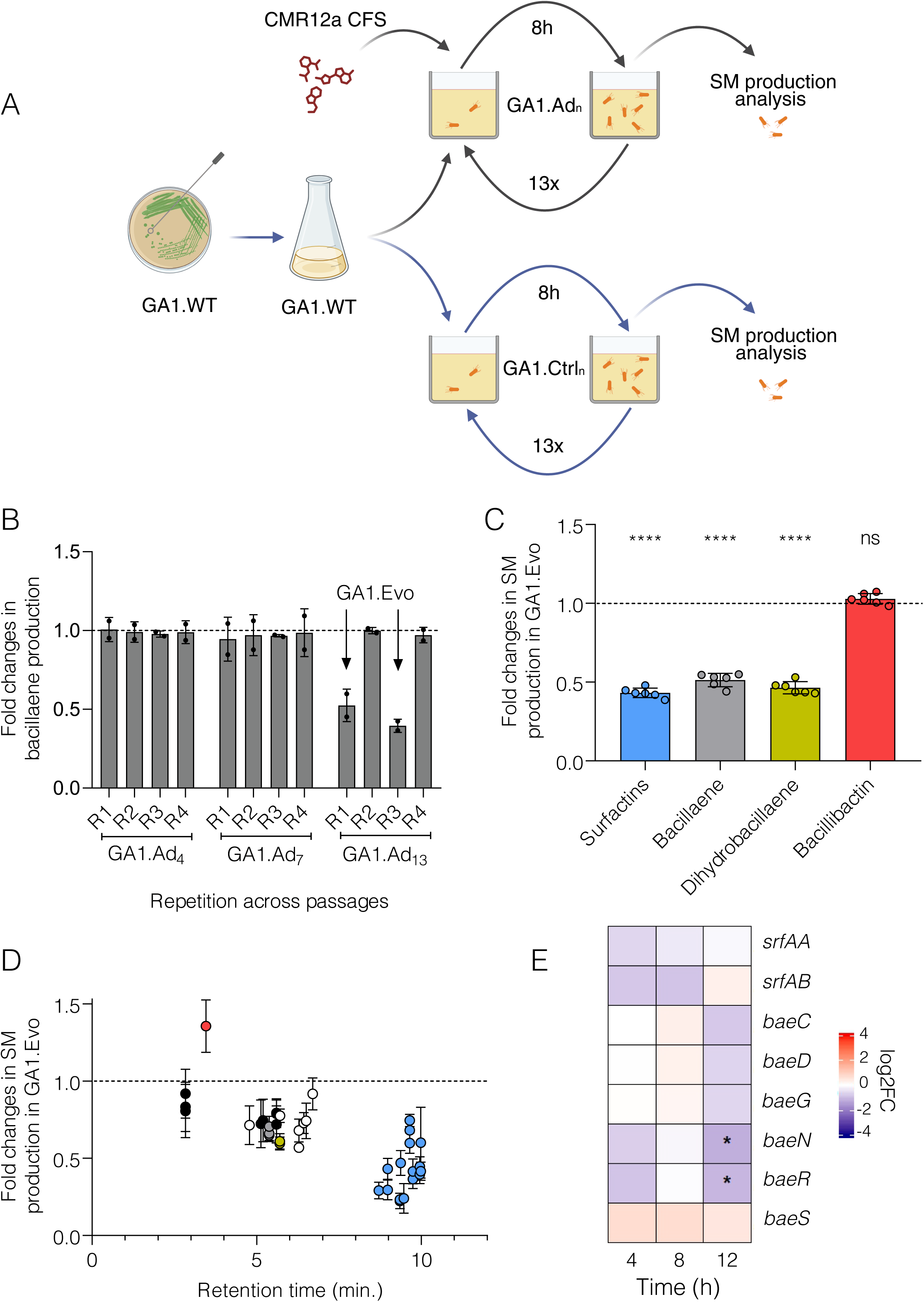
Repeated exposure to CMR12a CFS promotes adaptive diversification in *B*. *velezensis* GA1. A. Overview of the experimental evolution workflow applied to wild-type *B. velezensis* GA1 (GA1.WT) over 13 consecutive 8-hour culture cycles in the presence of *P. sessilinigenes* CMR12a cell-free supernatant (CFS). Each cycle consisted of growth in root exudate–mimicking medium [29] followed by transfer into fresh medium supplemented with CFS. Two independent lineages were established: GA1.Ad evolved in the presence of CFS and GA1.Ctrl grown without CFS. At the end of each cycle, cells were washed and either transferred to initiate the next passage or stored for downstream analyses. SM produced during each cycle were collected and analyzed by UPLC-MS (Created with BioRender.com). B. Relative SM production by *B. velezensis* quantified at three interaction passages (Ad_4_, Ad_7_ and Ad_13_). Bacillaene (used as representative SM) levels are expressed as fold changes of normalized peak areas relative to the untreated GA1 cells used as control condition (set to 1, horizontal black line). Data represent mean ± SD obtained from cultivation of four different isogenic colonies isolated on plates (R1 to R4), each repeated twice (n = 8). C. Relative SM production by the evolved lineage isolated from GA1 subpopulation occurring at passage 13. Metabolites were analyzed after 16 h of cultivation and production level is expressed as described in B. Data represent mean ± SD from three independent biological experiments, each performed with two technical replicates (n = 6). Statistical analysis was performed using an unpaired Welch’s t-test. Statistical significance: p < 0.0001 (****). D. Untargeted metabolomics performed for GA1.Evo after 16 h of cultivation. Each dot represents a detected feature (variants or adducts). Black dots correspond to uncharacterized metabolites, blue dots to surfactin-related molecular ions (see M&M), grey dots to bacillaene, green dots to dihydrobacillaene, red dots to bacillibactin, and white dots to other known SM families. Metabolite production is expressed as described in B. Data represent mean ± SD from three independent biological experiments, each performed with two technical replicates (n = 6). E. Differential expression of genes related to SM synthesis in the evolved lineage compared to WT as revealed by RNA-seq performed at different time points (4, 8 and 12 h). Red blocs indicate upregulated genes whereas blue blocs indicate downregulated genes expression. Differentially expressed genes (log_2_FC ≥ 1 or ≤−1 adj. p ≤ 0.05) obtained with PyDESeq2 are indicated with an asterisk (*).

We first analyzed SM production by cells from the two lineages collected after each batch and observed that, when exposed to *Pseudomonas* CFS, GA1.Ad consistently retained its ability to enhance the synthesis of (dihydro)bacillaene across the first 13 passages, indicating that sensing of metabolites from its competitor was maintained (Figure S2).

We next determined SM production by the GA1.Ad lineage at each passage compared to GA1.WT but without adding CMR12a CFS to test possible adaptation. No variation was detected in SM production across the first passages (Figure 1b and Figure S3) but the production rate of surfactin, bacillaene and dihydrobacillaene by GA1.Ad cells was consistently downregulated from subculture 13 and onward. As reported in some previous EE studies involving closely related species, we concomitantly observed the emergence of phenotypically distinct *B*. *velezensis* variants forming the dominant population starting from passage 13. Isogenic clones from this adapted population were isolated by plating and displayed a distinct morphology with larger colonies on agar LB compared with the ancestor.

This evolved GA1 lineage (hereafter defines as GA1.Evo) isolated from passage 13 was further analyzed for SM production. Targeted UPLC-MS analysis revealed significantly lower amounts of surfactin, bacillaene and dihydrobacillaene as compared to GA1.WT (Figure 1c). In contrast, the production of the siderophore bacillibactin remained unchanged in the evolved isolate suggesting that iron acquisition represents a non-negotiable fitness trait in this evolutionary context. Through untargeted metabolomics, we detected additional metabolite features with lower abundance in GA1.Evo (Figure 1d). Although the corresponding compounds remain to be characterized, this suggest that *B. velezensis* may adapt with decreased metabolite production beyond known SM.

CLPs and polyketides belong to the *B. velezensis* core metabolome and are non-ribosomal compounds synthesized by mega-enzymes encoded by giant biosynthetic gene clusters [42,43]. To evaluate if the observed lower SM production correlated with lower expression of the corresponding biosynthetic gene clusters (BGCs) in GA1.Evo, we performed RNA-seq analysis at different time points and manually inspected the differentially expressed genes (DEGs) (Figure 1e). In GA1.Evo, the genes encoding for the surfactin (*srfA*) operon were downregulated at 4 h and 8 h in agreement with the reduced production of the lipopeptide. In contrast, while most *bae* genes belonging to the (dihydro)bacillaene operon were not differentially expressed at 4 h and 8 h, we observed a global downregulation at 12 h. This late downregulation is in accordance with the reduced (dihydro)bacillaene production observed in GA1.Evo.

### Adaptation of GA1.Evo leads to better resource exploitation

Beyond SM-related transcriptional changes, global analysis of time-course RNA-seq data revealed extensive transcriptional reprogramming in GA1.Evo. The number of differentially expressed genes (DEGs) progressively increased over time to reach approx. 1000 DEGs at 12 h (Figure S4). To understand the biological function of the DEG, gene ontology (GO) enrichment analysis revealed temporarily distinct responses. At early time points, we observed enriched GO terms primarily associated with nucleotide metabolism, including ribonucleoside biosynthesis, purine nucleobase metabolism, and *de novo* IMP biosynthesis (Figure 2a). At later stages (8 h and 12 h), enrichment shifted toward amino acid metabolism, including pathways related to glutamine family, arginine–ornithine–histidine, and aromatic amino acid biosynthesis. In parallel, several amino acid biosynthetic pathways (e.g., pyruvate family, branched-chain amino acids, and methionine metabolism) were associated with downregulated genes. Multiple GO terms related to energy metabolism were also differentially regulated, with upregulated pathways including phosphoenolpyruvate-dependent sugar phosphotransferase systems, polysaccharide biosynthesis, iron–sulfur cluster assembly, and oxoacid metabolism, while others such as phosphate ion transport and inositol catabolism were downregulated (Figure 2a).

**Figure 2:**
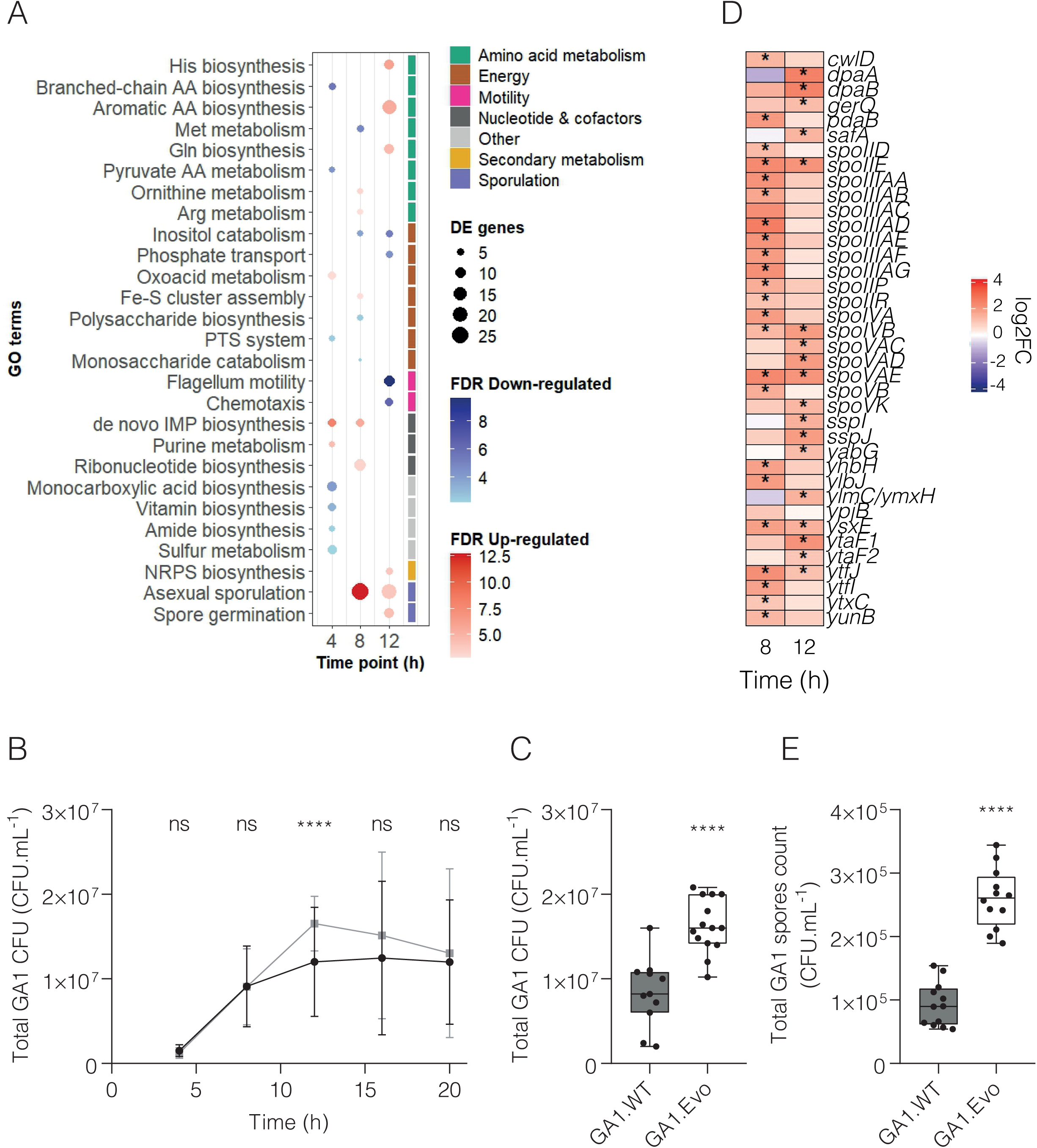
Adaptive evolution in *B. velezensis* GA1 drives extensive transcriptional reprogramming leading to increased biomass and sporulation. A. Gene ontology (GO) enrichment analysis related to biological processes performed on up- or downregulated genes with top GO R-package. RNA-sequencing was performed in evolved and wild-type GA1 at two different time points (8 and 12 h). Genes featuring a log2 fold change (FC) ≥ 1 or ≤−1 as compared to wild type GA1 (adj. p ≤ 0.05) were considered as differentially expressed. Red and blue circles indicate GO enrichment of up- or downregulated genes, respectively. Circle size represents the number of DE genes in each category. B. Growth kinetics of GA1 wild-type (GA1.WT) and evolved (GA1.Evo) populations. Total colony-forming units (CFU mL⁻¹), including both vegetative cells and spores, were quantified after 4, 8, 12, 16, and 20 h of cultivation. Data points represent the mean, and error bars indicate the SD. Data were obtained from three independent biological experiments, each performed with four technical replicates (n = 12). The black line corresponds to GA1.WT, whereas the grey line corresponds to GA1.Evo. Statistical comparisons between GA1.WT and GA1.Evo at each time point were performed using an unpaired Welch’s t-test. Statistical significance: p < 0.0001 (****). C. Total populations (CFU mL⁻¹) of GA1.WT and GA1.Evo isolates after 12 h of cultivation. Box plots display the median and interquartile range, with whiskers extending from the minimum to the maximum observed values. Data were obtained from three independent biological experiments, each performed with four technical replicates (n = 12). Statistical comparisons between GA1.WT and GA1.Evo were performed using an unpaired Welch’s t-test. Statistical significance: p < 0.0001 (****). D. Differential expression of genes related to sporulation in the evolved lineage compared to WT as revealed by RNA-seq performed at different time points (4, 8 and 12 h). Red blocs indicate upregulated genes whereas blue blocs indicate downregulated genes expression. Differentially expressed genes (log_2_FC ≥ 1 or ≤−1 adj. p ≤ 0.05) obtained with PyDESeq2 are indicated with an asterisk (*). E. Total spore counts of GA1.WT and GA1.Evo populations (CFU mL⁻¹) determined after 16 h of cultivation. Box plots elements are as described in panel C. Data were obtained from three independent biological experiments, each performed with four technical replicates (n = 12). Statistical comparisons between GA1.WT and GA1.Evo were performed using an unpaired Welch’s t-test. Statistical significance is indicated as follows: p < 0.0001 (****).

Many of these functional GO categories are closely linked to bacterial growth or physiology and we next investigated whether GA1.Evo exhibited specific growth dynamics compared to the wild type. We monitored growth kinetics over 20 h in liquid Re ½ medium under static conditions, and total cell abundance was quantified at five time points (4 h, 8 h, 12 h, 16 h, and 20 h). No significant differences in total cell numbers were observed between GA1.Evo and GA1.WT at 4 h and 8 h (Figure 2b) but the evolved strain displayed a significantly higher growth at 12 h (Figure 2c). As this biomass increase is associated with lower production of SM (Figure S5), we assume that it may derive from the fact that synthesis of these non-ribosomal compounds can be energetically costly as it involves a sophisticated mega-enzyme machinery using amino acids from the cellular pool as building blocks [44]. GO terms enriched in RNA-seq data indicate that central metabolism may be re-directed for more efficient growth. From 16 h onwards, both strains reached comparable population sizes. This convergence is likely due to nutrient depletion in the culture medium, limiting further growth.

Sporulation-associated GO terms, including spore germination (GO:0009847) and asexual sporulation (GO:0030436), were strongly upregulated at 8 h and 12 h, indicating a pivotal adaptive response. Further manual inspection of DEGs highlighted key genes involved in dipicolinic acid synthesis (*dpaA* and *dpaB*), genes encoding spore coat components (*cwlD*, *gerQ*, *safA*), multiple sporulation stage regulators (*spoII*, *spoIII*, *spoIV*, and *spoV*), and small acid-soluble spore protein (SASP) genes (*ssp*), which encode DNA-binding proteins essential for spore resistance (Figure 2d). We therefore quantified spore abundance at the same five time points used for total biomass assessment (4 h, 8 h, 12 h, 16 h, and 20 h). Although no differences were observed at the three first time points (Figure S6), spore population formed by GA1.Evo at 16h was significant enhanced compared to the wild type (Figure 2e). This is consistent with the enhanced growth observed at 12 h and may reflect faster nutrient consumption by GA1.Evo, enabling the population to reach stationary phase and initiate sporulation earlier than GA1.WT. Increased sporulation may also be induced by stress due to stronger nutritional limitation since the optimal biomass reached by GA1.Evo is higher.

### GA1.Evo can swarm but can’t swim

A closer look at the GO enrichment analysis also revealed strong downregulation of genes in GO-terms associated with bacterial motility in GA1.Evo such as those involved in flagellar assembly (GO:0071973) and chemotaxis (GO:0006935) (Figure 2a). In *B. velezensis*, bacterial movement mainly relies on two flagella-driven modes: swimming in liquid environments and swarming over semi-solid surfaces. While swimming primarily depends on the rotation of individual flagella that propel cells through liquid phases, swarming represents a collective movement that additionally relies on cellular differentiation and the production of surfactant molecules facilitating surface spreading [45,46].

Closer inspection of DEGs revealed significant downregulation at 12 h of several genes associated with the *fli* operon (Figure 3a), a key component of the flagellar machinery in *Bacillus* species [47]. Under swimming conditions (0.3 % m/v agar), we observed a strongly reduced motility of GA1.Evo and this difference became more pronounced over time, with the surface colonized by GA1.WT being more than 20 times larger than for GA1.Evo after 12 h (Figure 3b). The swarming behaviour was next evaluated by inoculating both strains on semi-solid medium (0.7 m/v agar) with colony expansion monitored over a 48 h period. An enhanced swarming ability of the evolved strain was observed with significantly larger surface spreading starting from 24 h (Figure 3c).

**Figure 3:**
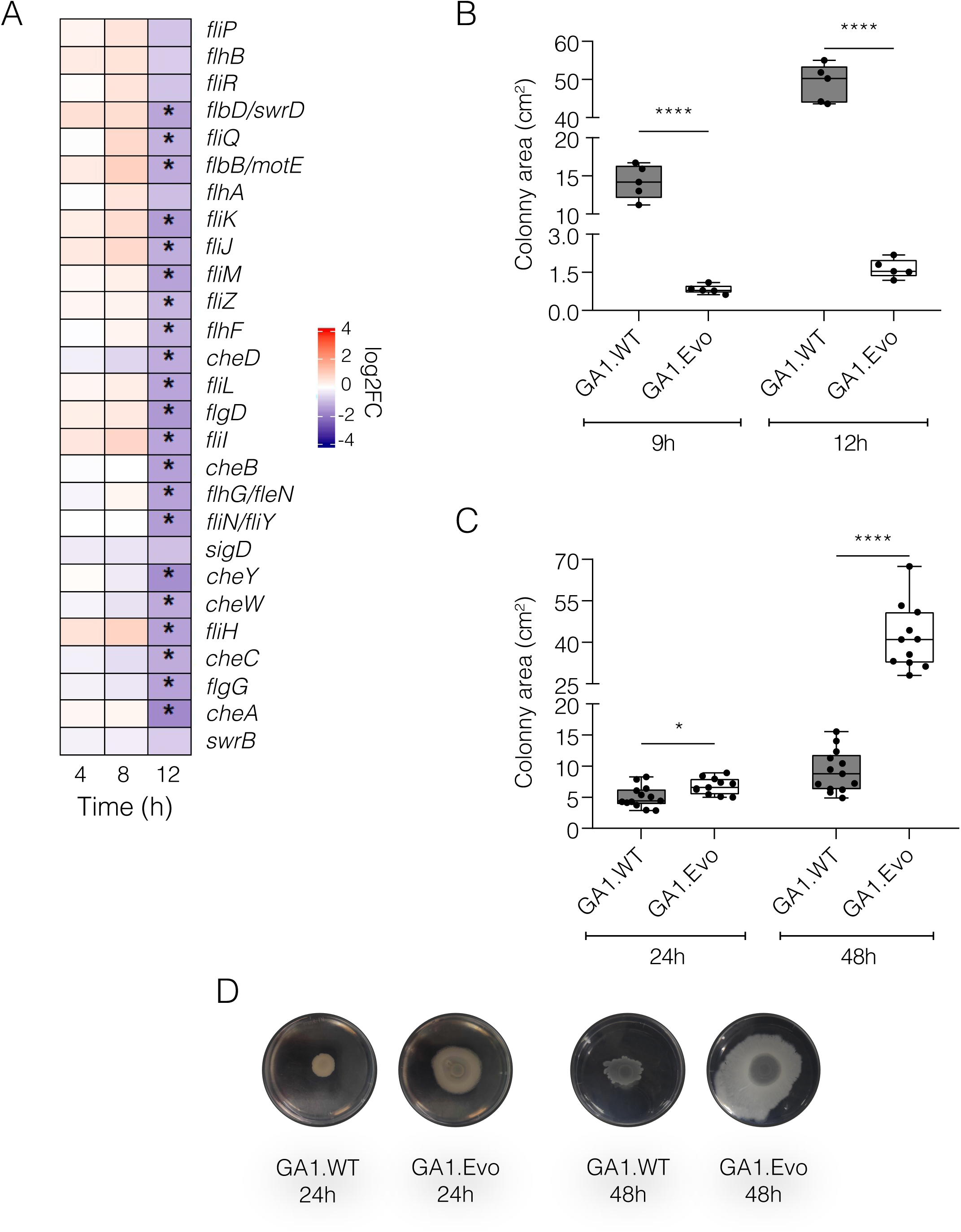
*B. velezensis* GA1.Evo has an improved swarming ability but a decreased swimming potential. A. Differential expression of genes related to motility in the evolved lineage compared to the wild type as revealed by RNA-seq performed at different time points (4, 8 and 12 h). Red blocs indicate upregulated genes whereas blue blocs indicate downregulated genes expression. Differentially expressed genes (log_2_FC ≥ 1 or ≤−1 adj. p ≤ 0.05) obtained with PyDESeq2 are indicated with an asterisk (*). B. Swimming ability of GA1.WT and GA1.Evo populations measured after 9 and 12 h of cultivation. Box plots display the median and interquartile range, with whiskers extending from the minimum to the maximum observed values. Data were obtained from one biological experiment, performed with five technical replicates (n = 5). Statistical comparisons between GA1.WT and GA1.Evo were performed using an unpaired Welch’s t-test. Statistical significance: p < 0.0001 (****). C. Swarming ability of GA1.WT and GA1.Evo populations measured after 24 and 48 h of cultivation. Box plots elements are as described in panel B. Data were obtained from two biological experiments, each performed with four technical replicates (n = 8). Statistical comparisons between GA1.WT and GA1.Evo were performed using an unpaired Welch’s t-test. Statistical significance: p < 0.05 (*) and p < 0.0001 (****). D. Representative pictures of the swarming ability of both GA1.WT and GA1.Evo taken after 24h and 48h of incubation at 30°C. Images are representative of at two biological replicates, each having at least four technical replicates.

### Decreased biofilm production by GA1.Evo is a conserved trait

Biofilm formation represents a key developmental trait for bacteria dwelling in competitive microbial environments. Biofilm matrix components include exopolysaccharides and structural proteins such as BslA, TasA, TapA and SipW, surrounded by a hydrophobic layer acting as a shield to protect cells from antimicrobial compounds and environmental stresses [5]. The close spatial organization within biofilms also promotes cooperative interactions and phenotypic heterogeneity, enabling cells to adopt distinct physiological states. This diversification reflects a bet-hedging strategy, whereby subpopulations specialize in functions such as matrix production, motility, or sporulation, increasing the likelihood that part of the population survives fluctuating and hostile conditions [48]. In this context, biofilm formation in *B. velezensis* may contribute to resilience and competitive fitness when exposed to metabolites produced by neighbouring microorganisms [49]. Therefore, we next investigated whether the evolved strain displayed altered biofilm formation.

Multiple genes in GA1.Evo associated with biofilm formation were strongly downregulated, notably genes belonging to the *eps* operon and other key matrix-associated genes including *bslA*, *tapA*, *tasA*, and *sipW* (Figure 4a). Downregulation was consistently observed across all time points, indicating a sustained transcriptional repression of biofilm matrix components in the evolved strain. This was supported by data from *in vitro* biofilm assays based on pellicle formation at the air–liquid interface. As shown in stereomicroscopy images acquired after 12 h, the evolved strain displayed a clearly altered and heterogeneous pellicle structure compared to GA1.WT (Figure 4b-c). More quantitatively, Crystal violet staining of matrix-embedded population confirmed the reduction in biofilm formation in the evolved strain (Figure 4d). Next, we used this quantitative analysis to determine whether such reduced biofilm-formation ability represented a stable trait. To this end, the evolved strain was sub-cultivated three consecutive times and Crystal violet staining was performed at each subculturing step. Interestingly, across all three sub-cultivations, the reduced biofilm phenotype remained consistent, indicating that the observed trait is maintained (Figure 4e). In parallel, we also monitored the level of SM production that also remained unchanged across the successive passages (Figure S7).

**Figure 4:**
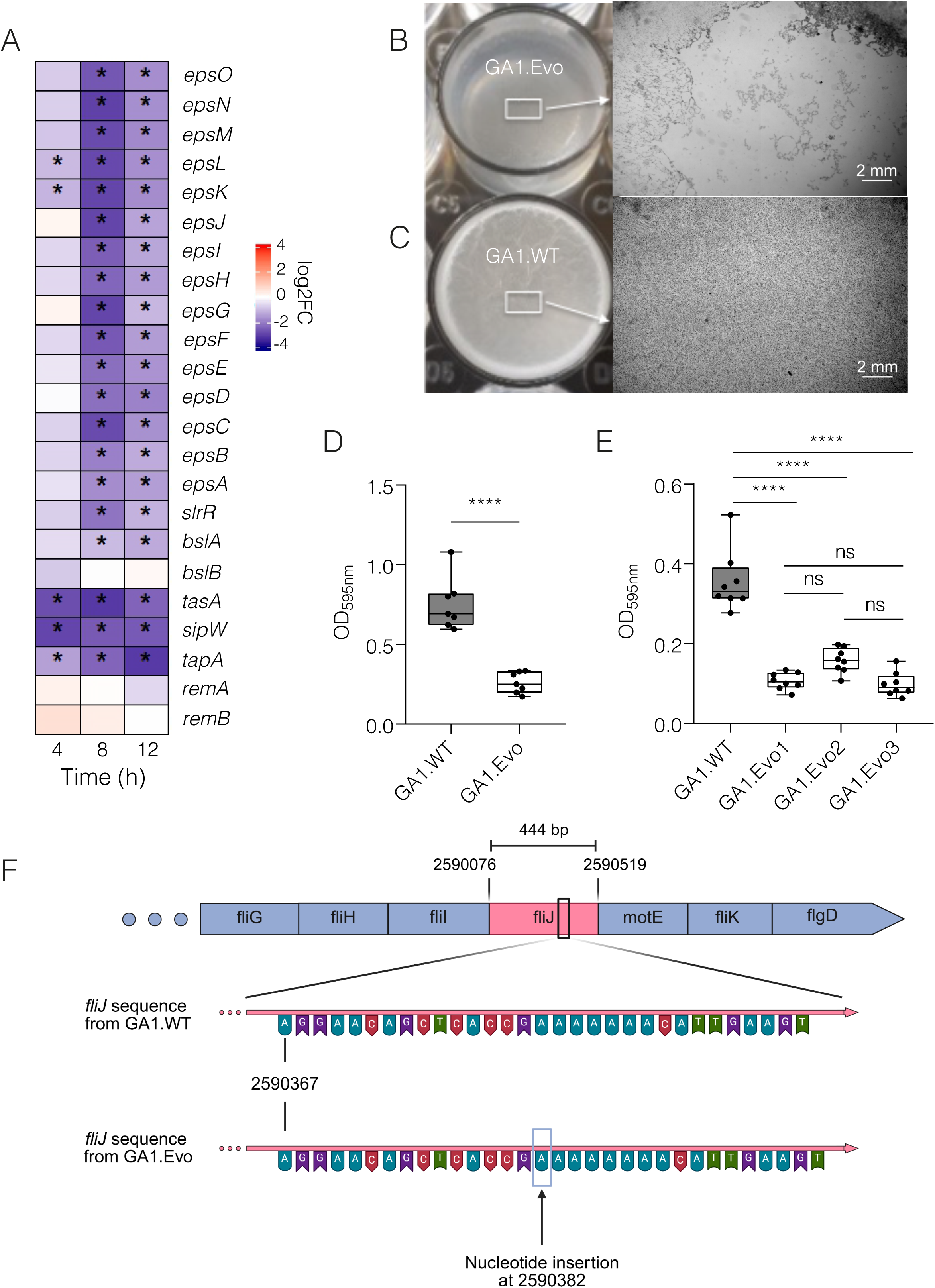
GA1.Evo biofilm formation ability is reduced and conserved over time. B. Differential expression of genes related to biofilm formation in the evolved lineage compared to the wild type as revealed by RNA-seq performed at different time points (4, 8 and 12 h). Red blocs indicate upregulated genes whereas blue blocs indicate downregulated genes expression. Differentially expressed genes (log_2_FC ≥ 1 or ≤−1 adj. p ≤ 0.05) obtained with PyDESeq2 are indicated with an asterisk (*). B-C. Representative pictures of biofilm formed by GA1.WT (B) and GA1.Evo (C) populations formed at the air-liquid interface after 12h of static incubation. Images were acquired with a stereomicroscope and are representative of at least three biological replicates. D. Assessment of biofilm formation at the air-liquid interface by GA1.WT and GA1.Evo after 12h of static incubation. Biofilm was quantified by crystal violet staining and measured spectrophotometrically at 595 nm (OD_595nm_). Box plots display the median and interquartile range, with whiskers extending from the minimum to the maximum observed values. Data were obtained from one biological experiment, performed with five technical replicates (n = 7). Statistical comparisons between GA1.WT and GA1.Evo were performed using an unpaired Welch’s t-test. Statistical significance: p < 0.0001 (****). E. Assessment of biofilm formation at the air-liquid interface by GA1.WT and GA1.Evo lineages after 12 h of static incubation. The evolved lineages were obtained after one (GA1.Evo1), two (GA1.Evo2), or three (GA1.Evo3) additional serial passages. Biofilm formation was quantified as described in D. Box plots elements are also as described in panel D. Data were obtained from one biological experiment with eight technical replicates (n = 8). Statistical comparisons between GA1.WT and the evolved lineages were performed using one-way ANOVA followed by Dunnett’s multiple comparisons test against the control condition (GA1.WT). Statistical significance: p < 0.0001 (****). F. Genomic organization of the chromosomal region surrounding *fliJ*, showing the neighbouring upstream and downstream genes, together with a nucleotide sequence comparison of the *fliJ* locus between GA1 wild-type and evolved strains. The evolved strain carries the insertion of a single adenine (A) nucleotide within the *fliJ* coding sequence, resulting in a frameshift mutation. Created with BioRender.com.

From these data, we hypothesized that genetic changes may have occurred during the evolutionary experiment that could explain the adaptive responses and their conservation. Whole-genome sequencing analysis of GA1.Evo identified a single mutation with a bp insertion occurring at 306 bp of the 444 bp *fliJ* gene (Figure 4f, Table S3). This gene encodes the soluble flagellar protein FliJ that forms part of the cytoplasmic ATPase complex (FliH–FliI–FliJ) of the type III secretion system involved in flagellar assembly [50]. In addition to its importance in flagellum export and, indirectly, its assembly, FliJ has been shown to interact with many other proteins in the *fli* operon [51–54]. Predictions made by the software PREDetector [55] highlighted the presence of 24 co-transcribed genes, all downstream of *fliJ* and involved in the mobility of *B. velezensis* (Table S3).

### GA1.Evo displays an increased competitiveness *in planta*

Since the adaptive response in GA1.Evo targets key developmental traits, we next investigated whether this may impact root colonization as one of the most desired traits for beneficial rhizobacteria. We therefore compared GA1.Evo with WT strain for early tomato root colonization capacity and *in planta* SM production.

After three days of incubation, GA1.Evo populations on tomato roots were significantly more abundant than those of the WT strain (Figure 5a). However, surfactin production rates were similar for both isolates as deduced from the amounts of CLP recovered from rhizosphere samples in these experiments, (Figure 5b). Considering the important role of surfactin in biofilm formation for *B. velezensis* GA1 [7,30], a similar production of this SM may suggest that biofilm formation is not impaired in GA1.Evo under *in planta* conditions.

**Figure 5:**
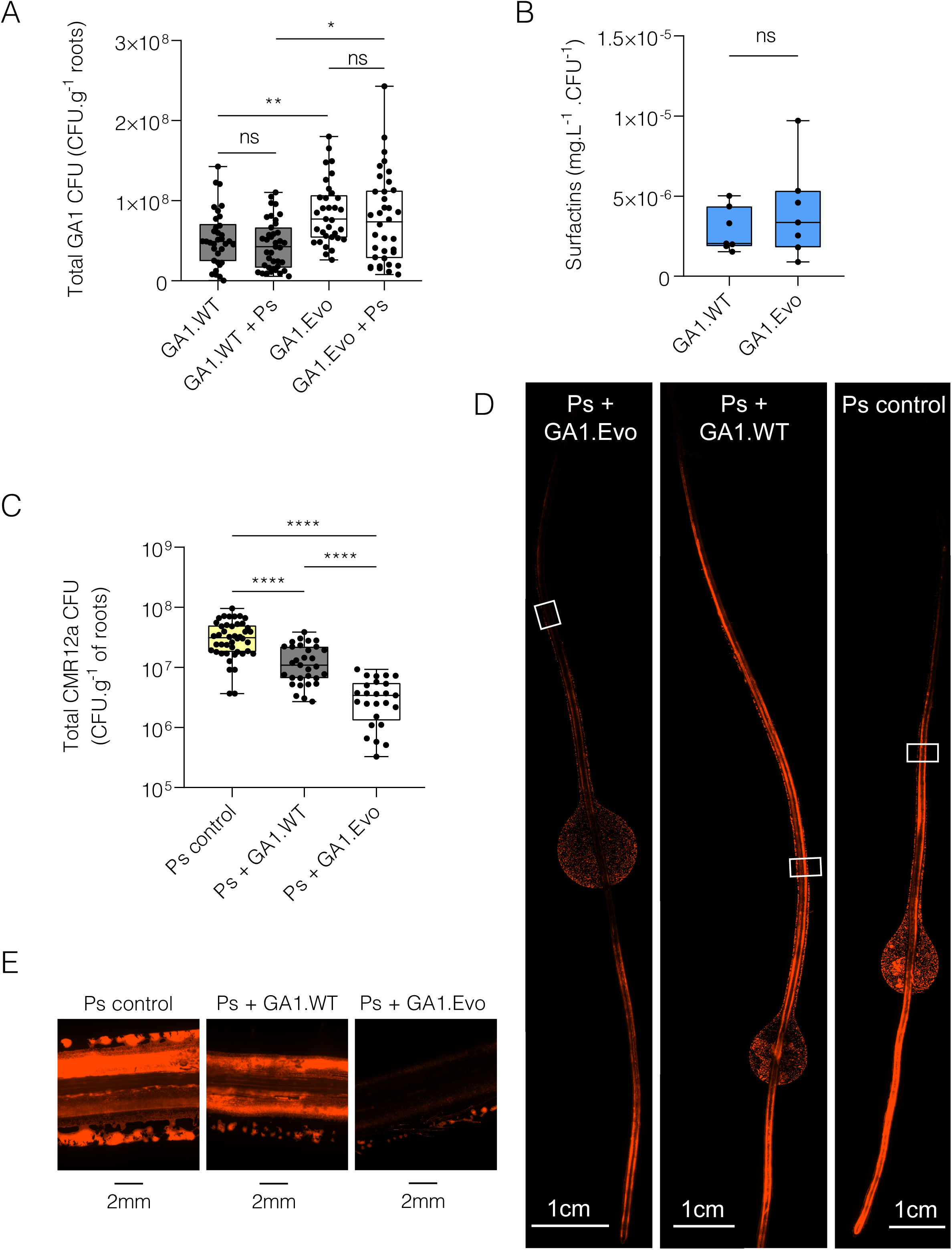
GA1.Evo effectively colonizes tomato roots and outcompete *Pseudomonas* competitor. A. Bacterial populations of GA1.WT and GA1.Evo recovered from roots of tomato plantlets three days post-inoculation. Two-week-old tomato plantlets were inoculated with either GA1.WT or GA1.Evo alone, or co-inoculated with *P. sessilinigenes* CMR12a (Ps), at a GA1 ratio of 95:5 (v/v). Box plots display the median and interquartile range, with whiskers extending from the minimum to the maximum observed values. Data were obtained from at least eight independent biological experiments, each performed with at least two technical replicates (n ≥ 20 per condition). Statistical comparisons were performed using Welch’s one-way ANOVA followed by Dunnett’s multiple comparisons test. Statistical significance: p < 0.05 (*) and p < 0.01 (**). B. Bacterial surfactin production by GA1.WT and GA1.Evo recovered from roots of tomato plantlets three days post-inoculation. Two-week old tomato roots were inoculated with either GA1.WT and GA1.Evo cells and surfactin was quantified three days post-inoculation. Box plot elements are as described in panel A. Data were obtained from at least five independent biological experiments (n ≥ 5 per condition). Statistical comparison between GA1.WT and GA1.Evo surfactin production was performed using an unpaired Welch’s t-test. C. Bacterial populations of *P*. *sessilinigenes* CMR12a (Ps) recovered from roots of tomato plantlets three days post-inoculation. Two-week-old tomato plantlets were inoculated with CMR12a alone or co-inoculated with either GA1.WT or GA1.Evo, at a GA1 ratio of 95:5 (v/v). Box plots elements are also as described in panel A. Data were obtained from at least eight independent biological experiments, each performed with at least two technical replicates (n ≥ 20 per condition). Statistical comparisons were performed using Welch’s one-way ANOVA followed by Dunnett’s multiple comparisons test. Statistical significance: p < 0.0001 (****). D-E. Representative epifluorescence images of *P. sessilinigenes* CMR12a::mCherry colonisation on tomato roots. Two-week-old tomato plantlets were inoculated with CMR12a alone or co-inoculated with either wild-type (GA1.WT) or evolved (GA1.Evo) B. *velezensis* GA1 cells. Images were acquired three days post-inoculation.

We next examined the impact of the presence of the *Pseudomonas* competitor CMR12a on root colonization ability of both GA1.Evo and WT. In these conditions, no significant impact of CMR12a on both GA1.Evo and WT root colonization was observed (Figure 5a). In contrast, we observed that *P. sessilinigenes* CMR12a colonized tomato roots less efficiently in the presence of *Bacillus*, with a more pronounced effect when co-inoculated with GA1.Evo (Figure 5c-d). This effect may be explained by a stronger spatial occupation by GA1.Evo, as observed during *in vitro* confrontations on agar plates (Figure S8).

## Discussion

In highly competitive environments such as the rhizosphere, plant-associated bacteria must continuously adapt to surrounding biotic interactions and signals. Here, we show that repeated exposure of *B. velezensis* to chemicals produced by a *Pseudomonas* competitor drives the emergence of a distinct subpopulation with specific traits. The evolved lineage notably displays enhanced root colonisation ability and higher competitive fitness upon co-inoculation compared with the wild-type strain. Efficient bacterial root invasion notably relies on swimming- and swarming-types of motility. While swimming motility is primarily required for chemotaxis allowing bacteria to reach the root surface, swarming motility is thought to play a more important role in root exploration and surface colonisation [56,57]. In our assays, bacteria were inoculated directly onto the root apex, thereby reducing the need for long-distance migration towards the plant. Under these conditions, the reduced swimming ability and the downregulation of chemotaxis-associated genes observed in GA1.Evo do not appear to impair its capacity to colonise roots. Furthermore, the semi-solid agarose medium used for the *in planta* assays is expected to favour swarming-type motility in *Bacillus*. The enhanced colonisation ability of GA1.Evo may therefore be partially linked to its increased swarming capacity compared with the wild-type strain.

Further insights into this altered motility phenotype were obtained from whole-genome sequencing, which revealed the presence of a single mutation in the evolved isolate corresponding to a one-nucleotide insertion in the *fliJ* gene. *fliJ* encodes a soluble flagellar protein belonging to the FliH–FliI–FliJ ATPase complex involved in type III flagellar export. This protein plays a central role in the transport of flagellar subunits and in the efficiency of flagellar assembly [51,58,59]. The frameshift mutation identified in GA1.Evo may result in a truncated FliJ protein. Interestingly, previous work in *Salmonella* demonstrated that the phenotypic consequences of *fliJ* mutations strongly depend on the mutation position within the gene [54]. Mutations located in the first half of the gene severely impaired swarming motility, whereas mutations occurring in the second part of the gene sequence may result in increased motility. Since the insertion identified in GA1.Evo occurs at position 306 of the 444 bp gene, our observations appear consistent with this latter scenario. Although further ultrastructural analyses are required to fully assess the consequences of the *fliJ* mutation, the reduced swimming capacity of the evolved lineage strongly suggest an impact on flagellum integrity and/or abundance since the single-nucleotide insertion identified in *fliJ* is expected to induce a frameshift mutation that may generate premature stop codons. Future investigations using transmission electron microscopy or fluorescent flagellar labelling would help determine whether the mutation affects flagellar assembly, morphology, or abundance.

Other studies have also used experimental evolution to investigate the adaptation of *Bacillus* spp. during repeated interactions with other organisms. EE performed on plant roots led to the emergence of distinct evolved subpopulations displaying improved adaptation to their host, resulting in enhanced colonization abilities [60,61]. Interestingly, several of these studies reported mutations affecting genes belonging to the *fli* operon or more generally involved in flagellar assembly and motility [60,62,63]. Although the mutation identified in the present study differs from those previously reported and does not affect the same gene, the recurrent occurrence of mutations within flagellar-associated genes may suggest that even if genetic adaptations are caused by stochastic changes, changes in the *fli* operon happen to be beneficial to the bacteria and are preserved. Whether this pattern reflects an increased susceptibility of this genomic region to mutation or, more likely, repeated positive selection acting on motility-related traits remains to be determined.

The altered motility phenotype of GA1.Evo may also indirectly affect biofilm formation. Under *in vitro* conditions, the evolved strain formed a less homogeneous pellicle at the air–liquid interface than the wild-type strain and this phenotype is associated with a reduced expression of genes involved in matrix production (*tasA*, *tapA*, *bslA*, and *eps*) [5,64] and with a lower production of surfactin, which are both known to contribute to biofilm formation in *B. velezensis* [30,65]. Reduced *in vitro* biofilm formation may additionally be linked to the altered swimming phenotype of GA1.Evo since pellicle formation first requires migration of planktonic cells towards the air–liquid interface [66,67]. However, whether this phenotype of reduced biofilm formation is maintained under plant-associated conditions remains unclear. Plant-derived compounds are known to strongly influence traits involved in root colonisation. In particular, plant polysaccharides can directly induce biofilm formation in *Bacillus* [68], while root exudates and pectin fragments stimulate surfactin production and modulate specialised metabolism in *B. velezensis* [7,30]. Consistently, the reduced surfactin production observed in GA1.Evo under *in vitro* conditions was no longer detected during root colonisation. This divergence may reflect the strong environmental dependency of SM biosynthesis. More generally, the pattern of bacterial secondary metabolites may vary considerably depending on growth conditions and lifestyle. Notably, secondary metabolite production has been reported to be strongly enhanced during biofilm-associated growth in plant-associated *Pseudomonas* spp. [69]. Together, these observations suggest that the reduced biofilm phenotype observed *in vitro* may not necessarily reflect biofilm formation under rhizosphere conditions. Further investigation of other SM produced by GA1.Evo *in planta* would help determining whether similar patterns are observed beyond surfactin. Further spatiotemporal investigation of biofilm formation by GA1.Evo *in planta* via quantification of biofilm-associated gene expression or visualization of biofilm matrix components on roots is necessary to better correlate enhanced colonisation capacity of GA1.Evo with altered biofilm formation in the rhizosphere.

Another hypothesis is that the improved root colonization ability of GA1.Evo may result from a reduced activation of plant immune responses. Our RNA-seq analyses revealed a decreased expression of several genes involved in flagellar assembly downstream of *fliJ*, suggesting a reduced production of the flagellar machinery. As flagellin is one of the major microbe-associated molecular patterns perceived by plants [70–72], a reduction in flagellar production could decrease host perception and facilitate bacterial establishment on the root surface. Consistent with this hypothesis, several plant-associated bacteria have been shown to downregulate flagellin expression after host colonization to limit flagellin-triggered immune responses [73]. Although the selective pressure applied during our EE is originated from bacterial competition and not the plant host, such potential immune evasion may arise as indirect consequences of adaptive changes affecting the flagellar machinery.

In addition to alter its growth dynamic, GA1.Evo also displayed changes in sporulation timing. Sporulation is a key adaptive trait allowing bacilli to persist under unfavourable biotic or abiotic conditions [74,75]. As bacterial populations reach high cell densities, *Bacillus* species often undergo developmental transitions leading to multicellular behaviours such as sporulation [76,77]. Although, in this study, early sporulation was observed only under *in vitro* conditions, it would be valuable to determine whether this phenotype is maintained *in planta* and whether it contributes to the persistence of the evolved strain in competitive root-associated environments. Moreover, further studies are needed to understand the link between the mutation observed in *fliJ* and the earlier sporulation of GA1.Evo.

Previous studies reported inhibitory effects of *Pseudomonas* spp. on *Bacillus* root colonisation [6,11,22]. However, under the experimental conditions used in this work, CMR12a did not negatively affect the colonisation of either *B. velezensis* strain. In contrast, both GA1 strains reduced CMR12a root colonisation, with a significantly stronger effect observed for GA1.Evo. Since surfactin production by *B. velezensis* is known to differ between *in vitro* and *in planta* conditions, this differential regulation may also extend to other secondary metabolites produced by GA1, including antibacterial polyketides. Nevertheless, direct inhibition of CMR12a by GA1-derived SM appears unlikely, as previous work showed that these metabolites do not impair CMR12a growth [11].

Rather than direct antibiosis, the enhanced competitive ability of GA1.Evo may primarily result from faster niche occupation and resource acquisition. This hypothesis is supported by our *in vitro* competition assays as well as the improved root colonisation capacity of GA1.Evo compared with the wild-type strain, both in the presence and absence of CMR12a. Consistent with this observation, GA1.Evo reached maximal biomass earlier than GA1.WT under *in vitro* conditions, suggesting a transcriptional reprogramming enabling more rapid resource utilisation. In competitive environments such as the rhizosphere, earlier nutrient consumption and faster establishment may provide a substantial ecological advantage by limiting resource availability for competing microorganisms.

While many studies have explored the co-evolution of bacteria with plants or fungi, this work provides new insights into the evolutionary impact of bacterium–bacterium interactions as mediated via chemical signalling without contact. It also further highlights the importance of considering long-term microbial interactions to implement our understanding of rhizosphere ecology in general and to better evaluate the adaptive potential of species such as *B*. *velezensis* as keystone biocontrol bacteria in particular.

## Acknowledgement

We gratefully thank Stéphanie Lambert (University of Liège) for her help in the construction of the GFP-tagged *B. velezensis* GA1.Evo strain, and Romain Thomas (University of Liège) for technical help in UPLC-MS. The authors Claude, Anthropic to assist with translation and language editing of portions of the manuscript. All content was reviewed and edited by the authors, who take full responsibility for the accuracy and integrity of the text.

## Author contributions

GB and AA performed most of the *in vitro* and *in planta* experiments. GB was involved in all metabolomics analyses. MZ and LVB performed the RNA-seq analyses, while SRM and SR carried out the genome sequencing analyses. GB, MZ, and MO primarily wrote the manuscript. All authors reviewed, commented on, and approved the final version of the manuscript. MO supervised the study and acquired funding.

## Conflicts of interest

The authors declare no conflicts of interest.

## Funding

G.B. and A.A. are recipient of a FRIA fellowship at the F.R.S.-FNRS (National Fund for Scientific Research in Belgium). S.R. and M.O. are respectively senior research associate and research director at the F.R.S.-FNRS. This work was supported by the projects PDR ID 40013634 and WEAVE ID T.0227.24/ G0AH724N from the FNRS/FWO.

## Data availability

All data generated or analyzed during this study are included in the paper and/or in the Supplementary files. Further information related to this article may be demanded from the corresponding authors G.B. or M.O.

## Notes

### Competing Interest Statement

The authors have declared no competing interest.

### Summary of Updates

Figures position adaptations + acknowledgement update

