## Supplementary Figures legends + Supplementary Tables for "*Bacillus* adaptation to *Pseudomonas* secondary metabolites enhances its root competitiveness"

### Supplementary Figures and Tables legends

***Figure S 1 :* Effect of *Pseudomonas* competitors' cell-free supernatants (CFS) on *Bacillus* spp. metabolites production*.***

Relative SM production by *B. velezensis* GA1 upon culture supplementation with *P. sessilinigenes* CMR12a, *P. defensor* WCS374 and *P. simiae* WCS417 CFS (4% v/v) after 8h of cultivation. SM productions are expressed as fold changes of normalized peak areas relative to the untreated GA1 cells used as control condition (set to 1, horizontal black line). Means and ± SD were calculated from data collected from three distinct experiments, each having three technical repetitions (n=9). Data were analyzed using one-way analysis of variance (ANOVA) when assumptions of homogeneity of variances were met. When variance heterogeneity was detected, Welch's ANOVA was applied instead. In both cases, mean values were compared to the control condition using Dunnett's multiple-comparison test adapted for unequal variances when appropriate. For the bacillibactin dataset, data were log10-transformed prior to statistical analysis to stabilize variance; graphical representations are shown using non-transformed values. Statistical significance is indicated as follows: $p < 0.05$ (*), $p < 0.01$ (**), $p < 0.001$ (***) and $p < 0.0001$ (****).

***Figure S 2 :* Evolution of *B. velezensis* secondary metabolites production upon *Pseudomonas* CFS culture supplementation*.***

Relative SM production by *B. velezensis* quantified at each of the 13 passages ($GA1.Ad_n$). Bacillaene (grey curve and points), and dihydrobacillaene (green curve and points) productions are expressed as fold changes of normalized peak areas relative to the untreated GA1 cells used as control condition (set to 1, horizontal black line). Means and ± SD were calculated from data collected from one experiment, having two technical repetitions (n=2).

**Figure S 3 : *B. velezensis* GA1 secondary metabolites production across three different serial passages.**

Relative SM production by *B. velezensis* quantified at three interaction passages ($Ad_4$, $Ad_7$ and $Ad_{13}$). Surfactins (blue), bacillibactin (red) and dihydrobacillaene (green) levels are expressed as fold changes of normalized peak areas relative to the untreated GA1 cells used as control condition (set to 1, horizontal black line). Data represent mean ± SD values obtained from four independent subpopulations (R1 to R4), each analyzed with two technicals replicates (n = 8).

***Figure S 4 :* Time course transcriptional reprogramming of *B. velezensis* GA1.Evo.**

Number of DEGs identified in GA1.Evo relative to the wild-type (GA1.WT) strain at 4, 8 and 12 h of cultivation. RNA-sequencing was performed on three independent biological replicate pools for each condition and time point. Genes exhibiting a log2 fold change (log2FC) ≥ 1 (upregulated; green) or ≤ −1 (downregulated; black) with an adjusted p-value ≤ 0.05 were considered differentially expressed.

***Figure S 5* : GA1.Evo spores counting over different cultivation times.**

Total spore counts (CFU $mL^{-1}$) of GA1.WT and GA1.Evo populations after 4 h – 8 h – 12 h of cultivation. Box plots display the median and interquartile range, with whiskers extending from the minimum to the maximum observed values. Data were obtained from three independent biological experiments, each performed with four technical replicates (n = 12). Statistical comparisons between GA1.WT and GA1.Evo were performed using an unpaired Welch's t-test.

***Figure S 6 :* GA1.Evo decreased BSMs production ability is a conserved trait.**

Relative SM production by the evolved lineage isolated from GA1 subpopulation occurring at passage 13 (GA1.Evo). SM were quantified after 12 h of static incubation. GA1.Evo was subjected to additional serial passages, yielding the successive lineages GA1.Evo1, GA1.Evo2, and GA1.Evo3 after one, two, and three additional passages, respectively. Surfactins (blue), bacillaene (grey) and dihydrobacillaene (green) levels are expressed as fold changes of normalized peak areas relative to the untreated GA1 cells used as control condition (set to 1, horizontal black line). Data represent mean and ± SD values obtained from one subpopulation, analyzed with two technicals replicates (n = 2).

***Figure S 7 : P. sessilinigenes* CMR12a colonies area during confrontation with both GA1.WT and GA1.Evo*.***

Evaluation of the impact of GA1.WT and GA1.Evo *on P. sessilinigenes* CMR12a colony growth. *CMR12a* colonies area were measured after 24 h of static incubation. CMR12a cells were either inoculated alone (left picture) or co-inoculated with either GA1.WT (central picture, CMR12a colony on the right of the picture), or with GA1.Evo (left picture, CMR12a colony on the right of the picture). Box plots display the median and interquartile range, with whiskers extending from the minimum to the maximum observed values. Data were obtained from three biological experiment, performed with at least one technical replicate (n ≥ 3). Data were analyzed using one-way ANOVA followed by Dunnett's multiple comparisons test against the control condition. Statistical significance: $p < 0.05$ (*), $p < 0.0001$ (****). Pictures are representative of the three biological replicates.

***Table S 1 :* Species and strains used in this study.**

| | **Relevant phenotype and description** | **References or sources** |
|---|---|---|
| ***Bacillus* strains** | | |
| *B. velezensis* GA1 | Wild Type | [1] |
| *B. velezensis* GA1 *ΔamyE*::*cat*-GFP (mut3.1) | GA1 disrupted of *amyE* gene. Chloremphenicol resistant, constitutive GFP reporter under Pveg promoter | This study |
| | **Relevant phenotype and description** | **References or sources** |
| ***Pseudomonas* strains** | | |
| *P. sessilinigenes* CMR12a | Wild Type | [2] |
| *P. sessilinigenes* CMR12a mCherry | mCherry tagged derivative of CMR12a | [3] |
| *P. defensor* WCS374 | Wild Type | [4] |
| *P. simiae* WCS417 | Wild Type | [5] |

***Table S 2 :*** **MZmine parameters used for detection of untargeted metabolite in *B. velezensis* GA1.Evo.**

| Step 1 | Parameter | | |
|---|---|---|---|
| **Mass detection** | MS level<br>Noize level | 1<br>100000 | |
| **Step 2** | **Parameter** | | |
| **ADAP chromatogram builder** | Mass List<br>Min group size in #scans<br>Group intensity threshold<br>Min highest intensity<br>m/z tolerance | masses<br>5<br>200000<br>500000<br>m/z | 0.005 ppm 20 |
| **Step 3** | **Parameter** | | |
| **Chromatogram deconvolution** | **Local minimum search**<br>Chromatographic threshold<br>Search min Rt range (min)<br>Min relative height<br>Min absolute height<br>Min ratio of peak top/edge<br>Peak duration range (min) | <br>85%<br>0.05<br>0%<br>500000<br>150%<br>0.05-1 | |
| **Step 4** | **Parameter** | | |
| **Isotope peak grouper** | m/z tolerance<br>Retention time<br>Monotonic shape<br>Maximum charge<br>Representative isotope | mz<br>0.02<br>ND<br>3<br>most intense | 0.001 ppm 5 |
| **Step 5** | **Parameter** | | |
| **Isotope peak finder** | Elements<br>m/z tolerance<br>Maximum charge | C, H, O, N, S, Cl<br>m/z<br>3 | <br>0.001 ppm 5 |
| **Step 6** | **Parameter** | | |
| **Join aligner** | m/z tolerance<br>Weight for m/z<br>Retention time tolerance<br>Weight for RT<br>Require same charge state<br>Compare isotope pattern | m/z<br>75<br>0.05 min<br>25<br>ND<br>ND | 0.002 ppm 20 |
| **Step 7** | **Parameter** | | |
| **Feature list filter** | minimum peaks in a row<br>minimum peaks in an isotope pattern<br>m/z<br>RT<br>Peak duration range<br>Chromatographic FWHM<br>Charge<br>Kendrick mass defect<br>Keep or remove rows | 3<br>2<br>ND<br>ND<br>ND<br>ND<br>ND<br>ND<br>Keep what match all criteria | |

***Table S 3*** **: Mutation found in *B. velezensis* GA1.Evo compared to the ancestral strain GA1.WT.**

| Reference | Contig | Position | Mutation type | Wild type | Mutation | Targeted gene |
|---|---|---|---|---|---|---|
| GCF_009734085.1 | NZ_CP046386.1 | 2590382 | Insertion | GAAAAAAA | GAAAAAAAA | *fliJ* |

| Gene ID | Refseq ID | Gene start position | Gene end position | Strand |
|---|---|---|---|---|
| GL331_RS13080 | WP_003154253.1 | 2590076 | 2590519 | + |

| Protein | GO_function | GO_process | Consequence of mutation | AA seq without mutation | AA seq with mutation |
|---|---|---|---|---|---|
| Flagellar export protein FliJ | Cytoskeletal motor activity | Bacterial-type flagellum-dependent cell motility | Introduction of multiple stop codons | MAYQFRFQKLLELKEN<br>EKDQTLTEYRQSVSEFET<br>VAEKLYENMSKKELLEKD<br>KESKLKCGMSVQEMRHYQ<br>QFVSNLENTIYHYQKLVIMK<br>RNEMNEKQEQLTEKNIEV<br>KKFEKMREKQFNMFALED<br>KAAEMREMDDISIKQFMIQGH* | MAYQFRFQKLLELK<br>ENEKDQTLTEYRQSV<br>SEFETVAEKLYENMS<br>KKELLEKDKESKLKC<br>GMSVQEMRHYQQFV<br>SNLENTIYHYQKLVI<br>MKRNEMNEKQEQLTEK<br>KH*SKEI*KNAGKTI*<br>YVRT*RQSCRDEGN<br>GRHFNQAVYDSGAL |
