## Supplementary figures and images for "*Bacillus* adaptation to *Pseudomonas* secondary metabolites enhances its root competitiveness"

### Figure S1

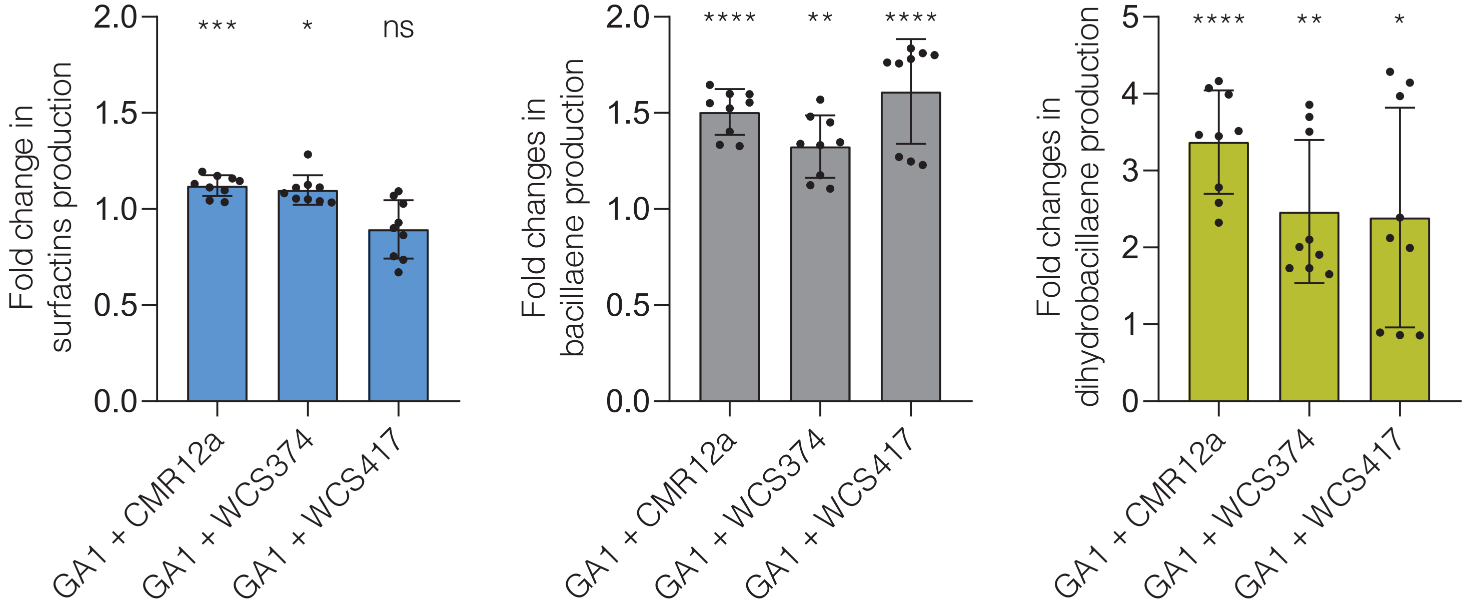

### Figure S2

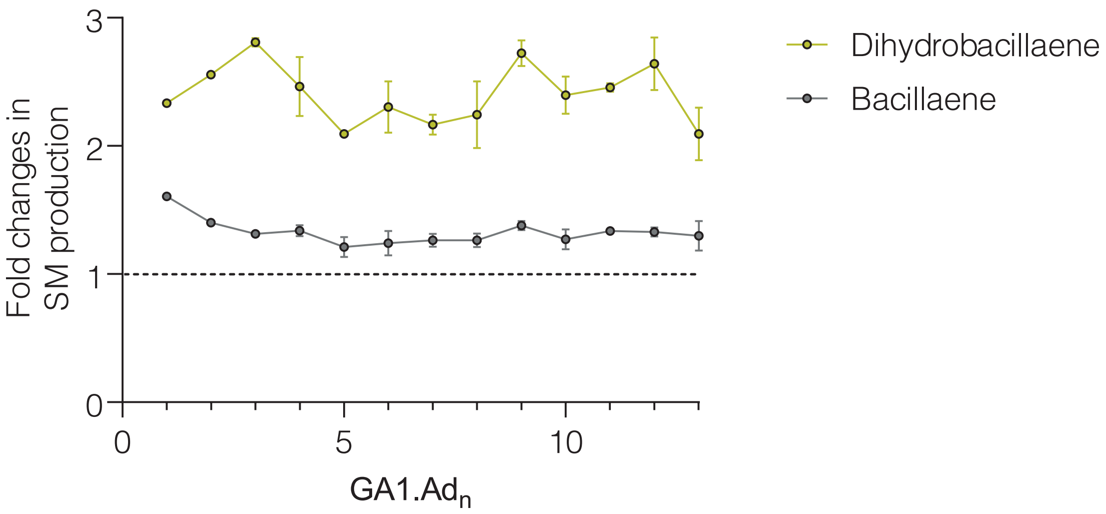

### Figure S3

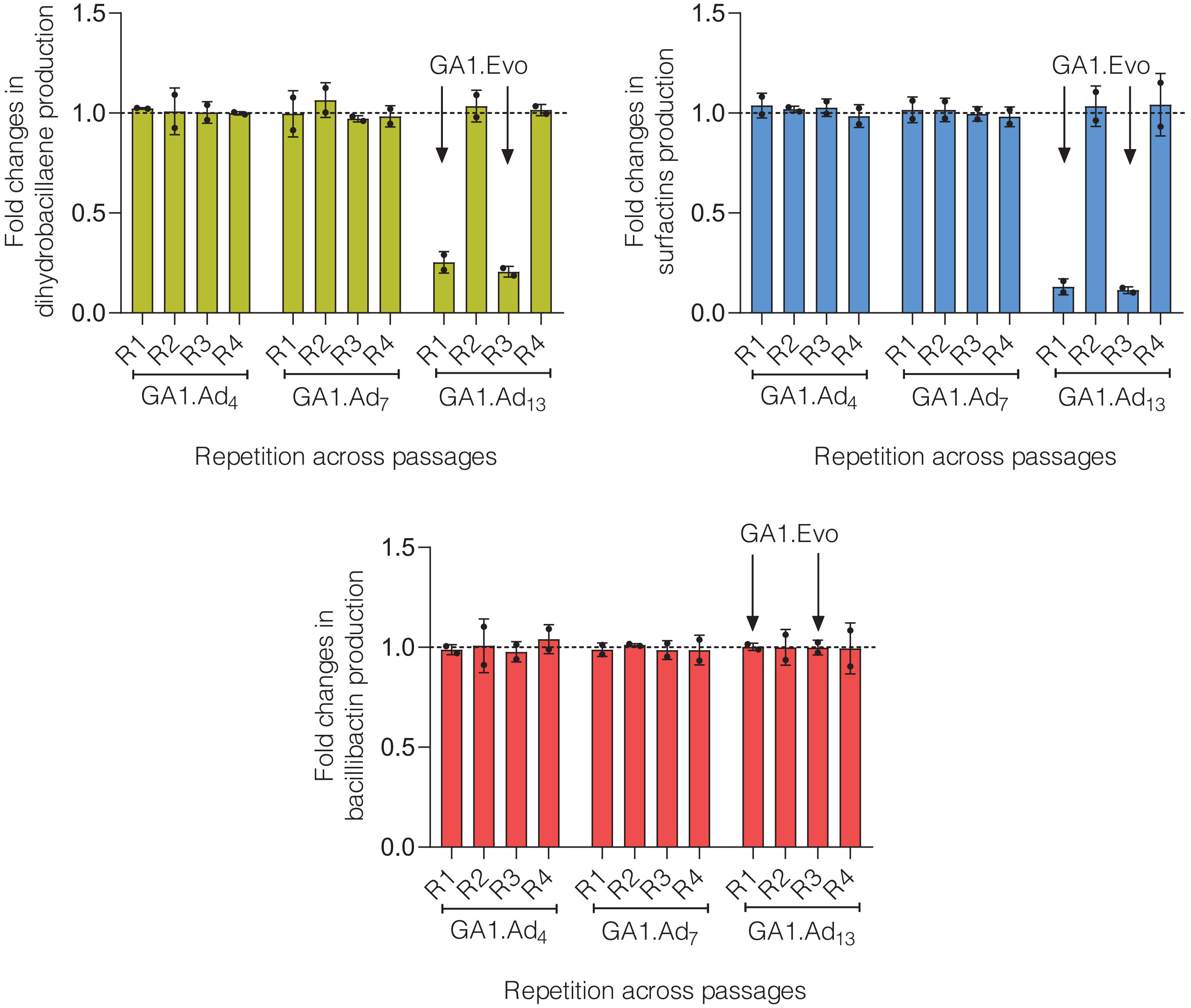

### Figure S4

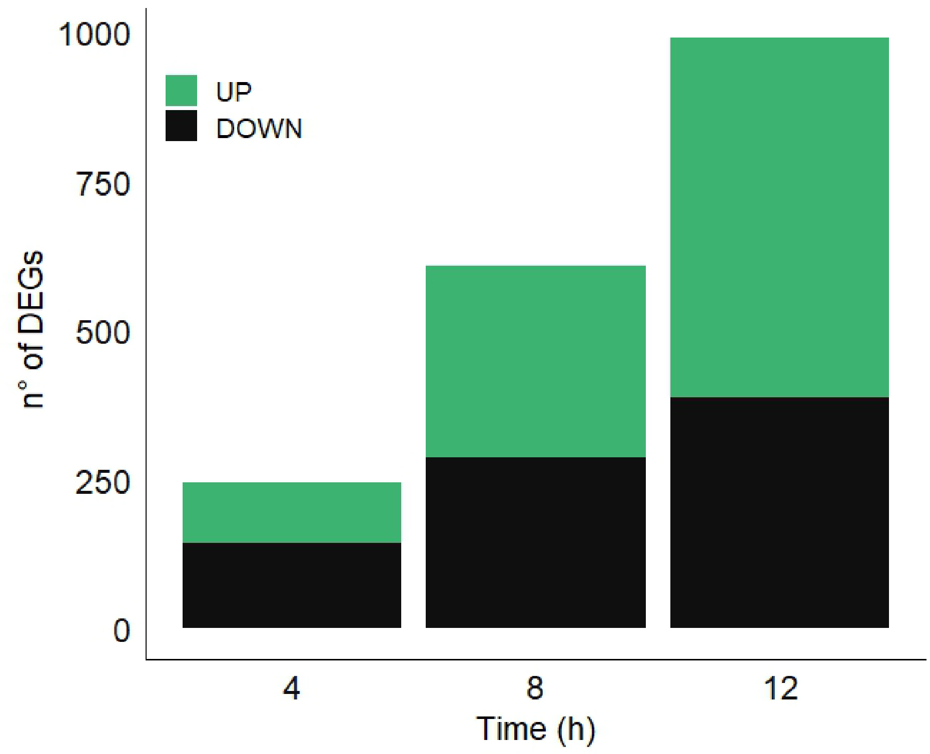

### Figure S5

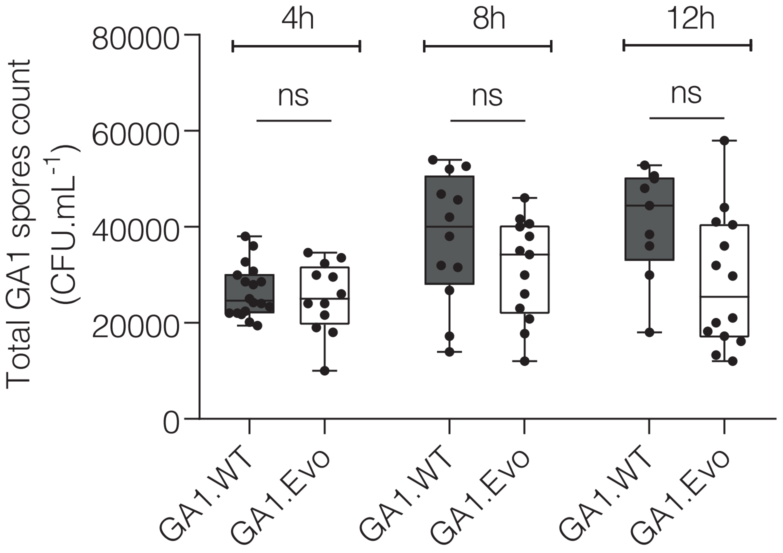

### Figure S6

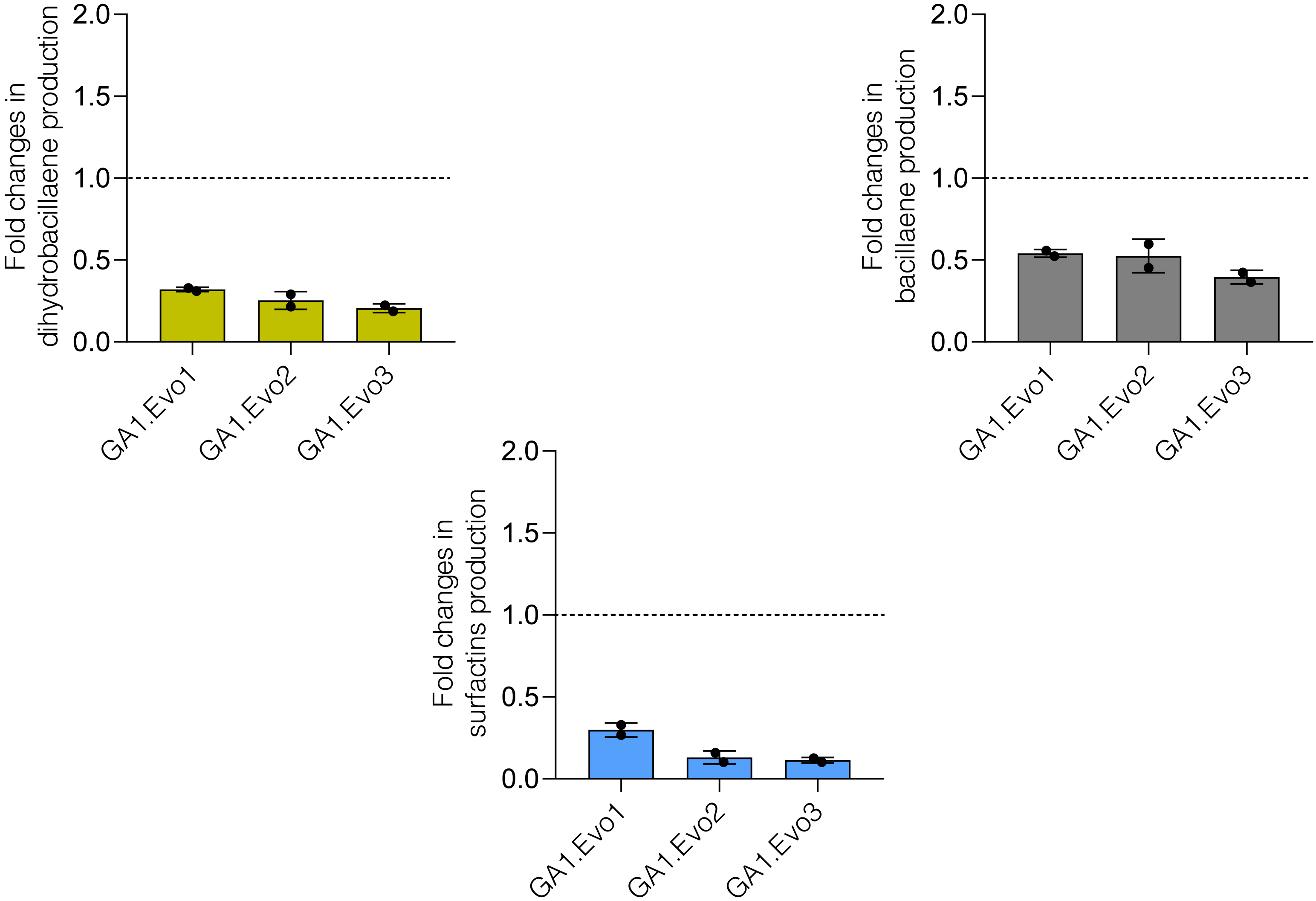

### Figure S7

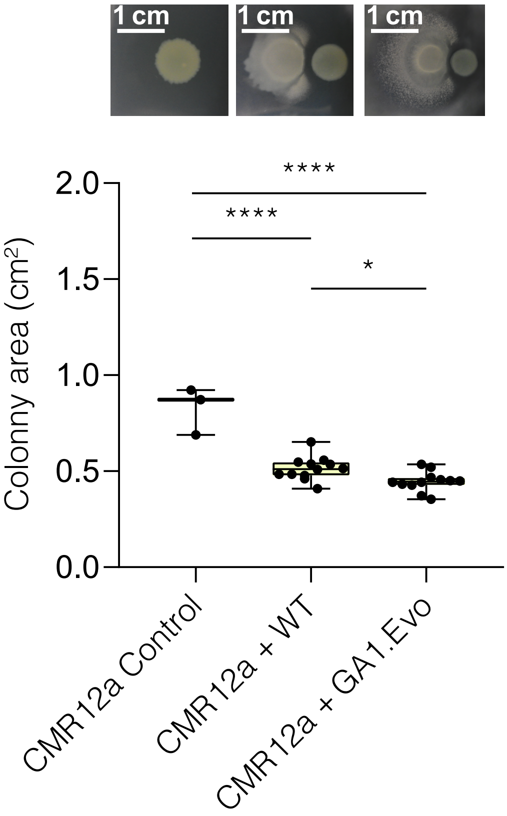
